# The naive T-cell receptor repertoire has an extremely broad distribution of clone sizes

**DOI:** 10.1101/691501

**Authors:** Peter C. de Greef, Theres Oakes, Bram Gerritsen, Mazlina Ismail, James M. Heather, Rutger Hermsen, Benjamin Chain, Rob J. de Boer

## Abstract

The human naive T-cell receptor (TCR) repertoire is extremely diverse and accurately estimating its distribution is challenging. We address this challenge by combining a quantitative sequencing protocol of TCRA and TCRB sequences with computational modelling. We observed the vast majority of TCR chains only once in our samples, confirming the enormous diversity of the naive repertoire. However, a substantial number of sequences were observed multiple times within samples, and we demonstrated that this is due to expression by many cells in the naive pool. We reason that *α* and *β* chains are frequently observed due to a combination of selective processes and summation over multiple clones expressing these chains. We test the contribution of both mechanisms by predicting samples from phenomenological and mechanistically modelled repertoire distributions. By comparing these with sequencing data, we show that frequently observed chains are likely to be derived from multiple clones. Still, a neutral model of T-cell homeostasis cannot account for the observed distributions. We conclude that the data are only compatible with distributions of many small clones in combination with a sufficient number of very large naive T-cell clones, the latter most likely as a result of peripheral selection.

## 1 Introduction

The human adaptive immune system employs a vast number (> 10^11^ [2]) of T lymphocytes, or T cells, to detect and dispose of pathogens. Most T cells express a single T-cell receptor (TCR) variant, which binds antigen in the form of a short peptide presented by the Major Histocompatibility Complex (pMHC) [3]. The TCR has to be specific to distinguish between self- and non-self-pMHC, but due to the large number of possible foreign antigens (> 20^9^) a specific TCR is nevertheless expected to bind many different pMHC (i.e., cross-reactivity) [29, 43]. The actual diversity of the TCR repertoire is unknown, but with improved sequencing techniques, estimates have risen by orders of magnitude from 10^6^ [1], 10^7^ [41], to over 10^8^ [39].

Generation of *αβ* T-cell diversity happens in the thymus, where thymocytes randomly rearrange gene segments to generate a TCR [36]. This heterodimer is generated by random recombination of Variable, Diversity, and Joining (V, D and J) segments for TCRB, and V and J segments for TCRA sequences [3]. Most variability arises due to random nucleotide insertions and deletions where the segments are joined [35]. Recent estimates of the potential number of TCRs produced by this V(D)J-recombination process range from > 10^20^ [54] to 10^61^ [33], which vastly outnumbers the number of distinct TCRs present in a human body. After generation of the TCR, T cells undergo positive and negative selection, which selects those T cells that have sufficient, but not too high, affinity for any self-pMHC [30]. About 3-5% of thymocytes survive selection [31] and enter the periphery as T cells that have not yet encountered foreign cognate antigen, i.e., as naive T cells.

How TCR diversity is maintained throughout life is an important question, as ‘gaps’ in the repertoire may allow pathogens to remain undetected [36, 53, 34]. The total number of CD4^+^ naive T cells stays relatively stable throughout life in the absence of cytomegalovirus (CMV) infection [51], while the CD8^+^ naive pool size declines substantially, irrespective of CMV status [51]. At the same time, thymic output of new T cells decreases because of thymic involution, making peripheral division of existing cells the main source of naive T cells from early adulthood onwards in humans [6, 22]. In the periphery, naive T cells compete for cytokines, such as IL-7, and need to interact with self-pMHC to survive [48, 47, 20]. Competition between T-cell specificities may reduce repertoire diversity when cells with some TCRs outcompete others, becoming more frequent [4]. Competitive T-cell dynamics lead to differences in the frequencies of TCRs, which determines the frequency distribution of their *α* and *β* chains. Hence, frequency distributions of TCR chains inform us about T-cell dynamics and how diverse repertoires are maintained.

To explain why some TCRs are more frequent in the naive repertoire than others, previous studies have mainly focused on fitness differences between T cells [46, 45, 17, 26, 8, 7, 9, 21]. Recently, the Mora and Walczak groups developed a probabilistic model that predicts the generation probability of any specific TCRA or TCRB sequence [35, 27]. They showed that these sequences (*σ*) differ several orders of magnitude in their probability 𝒫(*σ*) of being produced by V(D)J recombination in the thymus. Therefore, differences in generation probabilities may be an important factor in determining the frequency of TCR chains (i.e., TCRA and TCRB sequences) in the naive T-cell pool. Since it is not possible to directly assess the TCR repertoire in its entirety, we study a mathematical model of neutral T-cell dynamics, as well as various phenomenological clone-size distributions. We also account for the sampling process, which is an inevitable aspect of all experimental measurements of the TCR repertoire. The combination of modeling and careful quantitative measurements of TCRA- and TCRB-sequence frequencies in the naive repertoire in peripheral blood from healthy humans was used to test which distributions are compatible with the experimental observations. Remarkably, we find that only distributions which consist of many small but also some large clones are compatible with the observed frequency distribution of TCR chains from naive T cells in blood samples.

## 2 Results

To study the frequency distribution of TCR chains in the naive T-cell compartment, we reanalyzed our TCRA- and TCRB-sequence data published in [37]. In brief, peripheral blood mononuclear cells (PBMCs) from two adult volunteers were FACS-sorted into naive (CD27^+^CD45RA^high^) and various memory CD4^+^ and CD8^+^ populations. *TCRA* and *TCRB* mRNA was reverse transcribed to cDNA molecules to which unique molecular identifiers (UMIs) were attached, followed by PCR-amplification and high-throughput sequencing (HTS) on an Illumina MiSeq platform. Sequence reads were processed using a customized version of the Decombinator pipeline [49], with an improved error correction on UMIs to more reliably estimate the frequency of TCR chains in the samples (SI 4.2). Additionally, we used the RTCR pipeline [15] for comparison (SI 4.2).

### 2.1 *α* and *β* chains present in both naive and memory subsets have high generation probabilities

Using the V(D)J-recombination model of Marcou *et al.* [27], we predicted the generation probabilities 𝒫(*σ*) of all TCRA and TCRB sequences in our datasets. As expected, we observed a wide range of 𝒫(*σ*) values, for *β* chains on average multiple orders of magnitude lower than *α* chains (due to their recombination of the D-segment). The generation probability distributions of sequences derived from naive and memory T cells appeared to be identical (Fig. 1A). Thus, our data provide no evidence that the V(D)J-recombination process is selected for producing chains that are more likely to become part of an immune response. Remarkably, the *α* and *β* chains observed both in memory and the corresponding naive samples, were strongly enriched for high 𝒫(*σ*) (Fig. 1A). Thus, although generation probabilities tend to be similar between naive and memory T cells, we find that *α* and *β* chains that occur in both have high 𝒫(*σ*).

**Figure 1:**
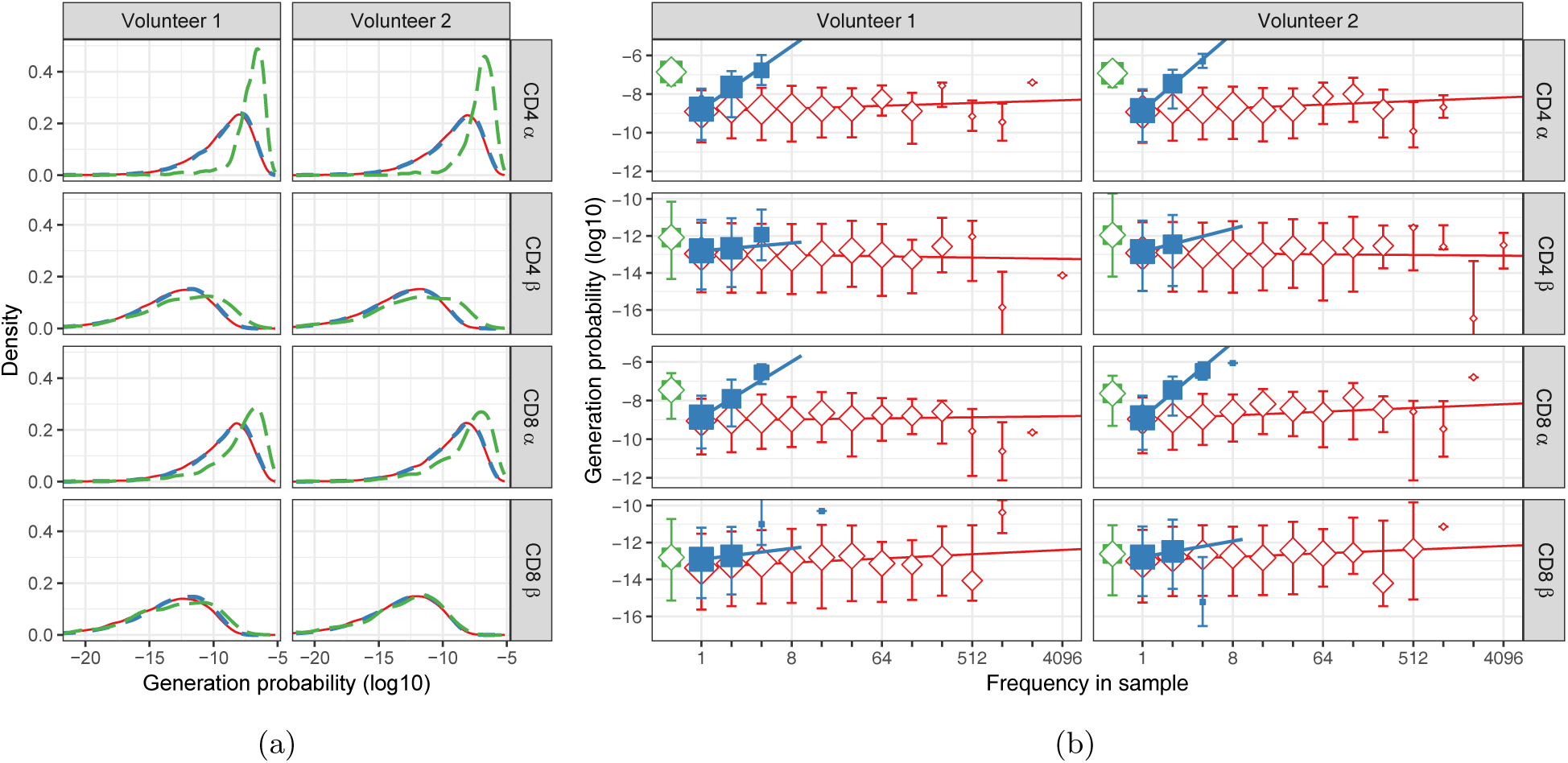
Generation probabilities of TCRA and TCRB sequences from naive and memory T cells. **A.** For each sequence *σ* in our dataset, the generation probability 𝒫(*σ*) was predicted using IGoR [27]. The distribution of generation probabilities (log10) for *α* and *β* chains from CD4^+^ and CD8^+^ from two volunteers is shown. Blue dashed: naive, red solid: memory, and green long-dashed: overlap (i.e., sequences observed in both naive and memory). **B.** The median 𝒫(*σ*) is shown for each observed frequency class (log2-bins) in naive (blue squares) and memory T-cell (red diamonds) samples. 𝒫(*σ*) of the overlapping chains is shown in green for reference (irrespective of frequency). Symbol sizes indicate numbers of sequences for each frequency class. Error bars represent the 25% and 75% quartiles, solid lines indicate linear regression between observed frequency and 𝒫(*σ*), weighted by the number of sequences with that frequency.

*α* and *β* chains can occur in both memory and naive samples for multiple reasons. The TCRA and TCRB sequences of (large) memory T-cell clones are expected to be observed in the naive sample due to impurities in the FACS-sorting. However, if the overlap were the result of contamination only, the 𝒫(*σ*) of the sequences would be expected to reflect those of the memory subsets. Since the overlap is markedly enriched for high generation probabilities, most of it cannot be caused by contamination. A better explanation arises because our sequencing data lacks information on *α* and *β* chain pairing. Overlapping chains in our data can be derived from multiple clones with the same *α* chain but different *β* chains, or vice versa, and chains that are most likely to be generated by V(D)J recombination (i.e., those with high 𝒫(*σ*)) are expected to occur in many clones. Thus, for chains with a high 𝒫(*σ*), there is a high probability that at least one of the many clones expressing the same *α* chain, but different *β* chains (or vice versa), has been part of an immune response. This can explain the observation that chains overlapping between naive and memory have high 𝒫(*σ*).

### 2.2 Frequently observed *α* and *β* chains in the naive samples tend to have high generation probabilities

Within the naive T-cell samples, the vast majority of TCRA and TCRB sequences were observed only once, and most frequencies fall within the range from 1 to 5. As expected, in the memory samples, which contain clonally expanded T cells, much more frequently observed sequences were present, with a substantial number of *α* and *β* chains observed more than 1000 times. The few sequences observed with a frequency higher than 5 in the naive samples occurred in almost all cases (94.6%) also in the corresponding memory subset. As discussed above, these could partly be explained by impurities in the FACS-sorting. Therefore, when correlating observed frequency and generation probabilities, we excluded from the analysis any sequence that overlapped between naive and memory subsets from an individual, in order to enrich the naive population for truly naive T cells. Note that this is a conservative approach, since it also excludes truly naive but frequent TCRA and TCRB sequences with high 𝒫(*σ*) (Fig. 1A).

The frequencies of sequences in all purified naive subsets show a positive correlation with the probability of being produced by V(D)J recombination (Fig. 1B: blue). In our naive T-cell samples, the median 𝒫(*σ*) of the *α* chains that were observed at least three times was about 154-fold higher than for those that have only been observed once (*p* < 10^−15^, Wilcoxon test). This trend was weaker for *β* sequences (∼2.5-fold, *p* < 0.01, Wilcoxon test), but still stronger than for memory subsets (1.65- and 1.03-fold for *α* and *β*, respectively). These results are consistent with the explanation that frequently observed chains in the naive compartment are derived from many clones because of their high generation probabilities. The most frequently observed chains from memory T cells, on the other hand, are derived from large *αβ*-clones, without enrichment for high generation probabilities (Fig. 1B: red).

### 2.3 Frequently observed TCR chains cannot be attributed only to multiple RNA molecules per cell

The frequency measurements are influenced both by the number of cells in the sample expressing a given chain, and by the number of *TCRA* and *TCRB* mRNA molecules per cell. Although we have determined the average number of mRNA molecules per cell [37], this distribution has a high variance, perhaps due to transcriptional bursting. From the full naive pool (∼10^11^ cells), the probability for a cell to be part of a sample of ∼10^6^ cells is very small (∼10^−5^). However, cells present in the sample can contribute multiple mRNA molecules with a probability that is likely to be much higher. In order to exclude this uncertainty in analyzing the frequency of *α* and *β* chains in the naive repertoire, we performed an additional experiment. We sorted naive T cells from an additional volunteer, and after sorting split the naive T cells into three subsamples before mRNA extraction. We then carried out library preparations and sequenced TCRA and TCRB sequences from each subsample. In this experiment, sequences observed in more than one subsample must have been derived from different cells, and cannot be a result of sequencing multiple mRNA molecules from a single cell. Repeated sequences are therefore direct evidence of frequent TCR chains.

In total 17199 (3.8%) TCRA sequences, and 5793 (0.71%) TCRB sequences, were observed in more than one subsample (Fig. 2A), firmly establishing the existence of a substantial number of frequent TCR *α* and *β* chains in the full naive repertoire. The frequently observed TCR *α* chains in our samples are dominated by sequences with high 𝒫(*σ*): the median generation probability of TCRA sequences observed in two and three subsamples was 56- and 165-fold higher, respectively, than those observed only once (Fig. 2B). This reconfirms the role of generation probabilities in the frequency of *α* chains in the naive TCR repertoire. We also observed substantial numbers of TCRB sequences in two and even three subsamples, but with a remarkably different 𝒫(*σ*) trend. While *β* chains observed in two subsamples are mildly enriched for high generation probabilities, those observed in three subsamples have hardly any enrichment for high 𝒫(*σ*) (Fig. 2B). Instead, their generation probabilities tend to be even lower than those of the sequences observed in two subsamples, and are more similar to the generation probabilities of TCRB sequences with incidence 1. Note that these trends were observed both with and without removing the sequences that were also observed in memory (Fig.2, grey versus colored bars).

**Figure 2:**
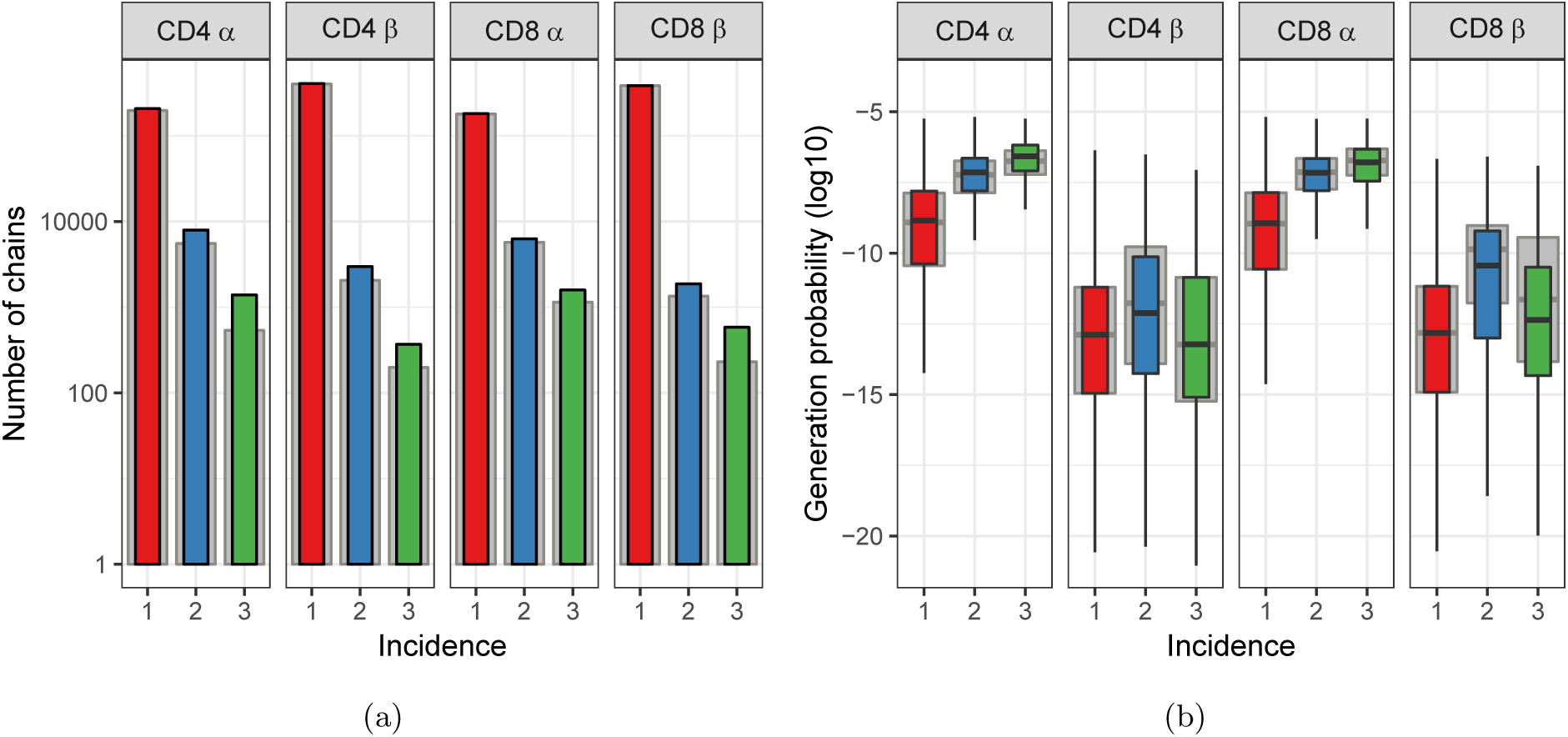
Subsampling naive T cells confirms that frequently observed TCRA but not TCRB sequences have high generation probabilities. **A.** The number of TCRA and TCRB sequences observed in 1, 2 or 3 subsamples. The grey background bars show the results after removing all sequences that were also observed in the corresponding memory samples. **B.** Generation probabilities 𝒫(*σ*) (log10) of chains observed in 1, 2 or 3 subsamples.

We also performed an orthogonal analysis to verify these observations independent of IGoR (SI 4.3). *α* or *β* chains that are likely products of the V(D)J-recombination process, and thus repeatedly generated, will also be more likely to occur in different individuals (i.e. be more “public”). We therefore measured sharing between those TCR chains observed in 1, 2, or 3 naive subsamples, with TCRA and TCRB repertoires sequenced from unfractionated blood samples collected from 28 healthy donors. Both *α* and *β* chains observed in two or three subsamples were found to be significantly more often shared with this independent cohort than those observed once (Fig. S3A). The most frequent TCRA sequences, which were seen in three subsamples, showed the highest sharing degree, consistent with their strongest enrichment for high generation probabilities. The relatively small number of most frequent TCRB sequences (i.e., those observed in three subsamples), did not show increased inter-individual sharing compared to the TCRB sequences observed in two subsamples. Additional comparison with publicly available TCRB data from a large cohort [13] showed, consistent with the trends in 𝒫(*σ*), that the most frequently observed *β* chains, with incidence 3, were less public than sequences observed in two subsamples (Fig. S3B). Thus, our findings surprisingly indicate that the probability to be generated by V(D)J recombination may explain the frequency of *α*, but not *β* chains, although both chains are derived from the same samples of cells.

### 2.4 A model of neutral naive T-cell dynamics is compatible with the observed frequency distribution of TCRA, but not TCRB sequences

Our aim is to study the distribution of naive T-cell clones, which are identified by the combination of their *α* and *β* chain. Our results indicate that the frequencies of chains in our samples are not only the result of true heterogeneity in clone sizes, but also by summation over multiple clones. Hence, one cannot directly tell the TCR distribution of naive T-cells from HTS samples of *α* and *β* chains. Instead, we use mathematical models to predict clone-size distributions and compare samples from these distributions with the HTS data of the subsample experiment (described above in Section 2.3). This approach allows us to include the different mechanisms determining TCRA and TCRB frequency, and study which distributions agree best with our data. Clones that for some reason are large in the pool are more likely to contribute cells with their *α* and *β* chains to the samples. At the same time, *α* and *β* chains with high generation probabilities will be expressed by many clones. Mathematical models allow us to take both these factors, and the stochastic nature of the sampling process, into account when predicting the frequency of *α* and *β* chains in samples.

We first develop a simple neutral model, similar to Hubbell’s Neutral Community Model [19], for the dynamics of clones (Fig. 3A). A neutral model assumes that the TCR of a naive T cell does not affect its lifespan or division rate. Consider a pool of *N* naive T cells, from which cells are removed by cell death or by priming with antigen, leading to differentiation into a memory population. A fraction *θ* of these cells is replaced by thymic production of new clones and the remaining fraction 1 − *θ* gets replaced by division of cells present in the pool. When simulating the naive T-cell pool with this model, the clone-size distribution approaches a “steady state” (not shown). We use this steady-state distribution, for which we have an analytical expression (SI Section 4.4) to predict the size of clones in the naive T-cell pool. As the contribution of thymic output decreases during aging [44, 52], we evaluate the model for a wide range of values for *θ*.

**Figure 3:**
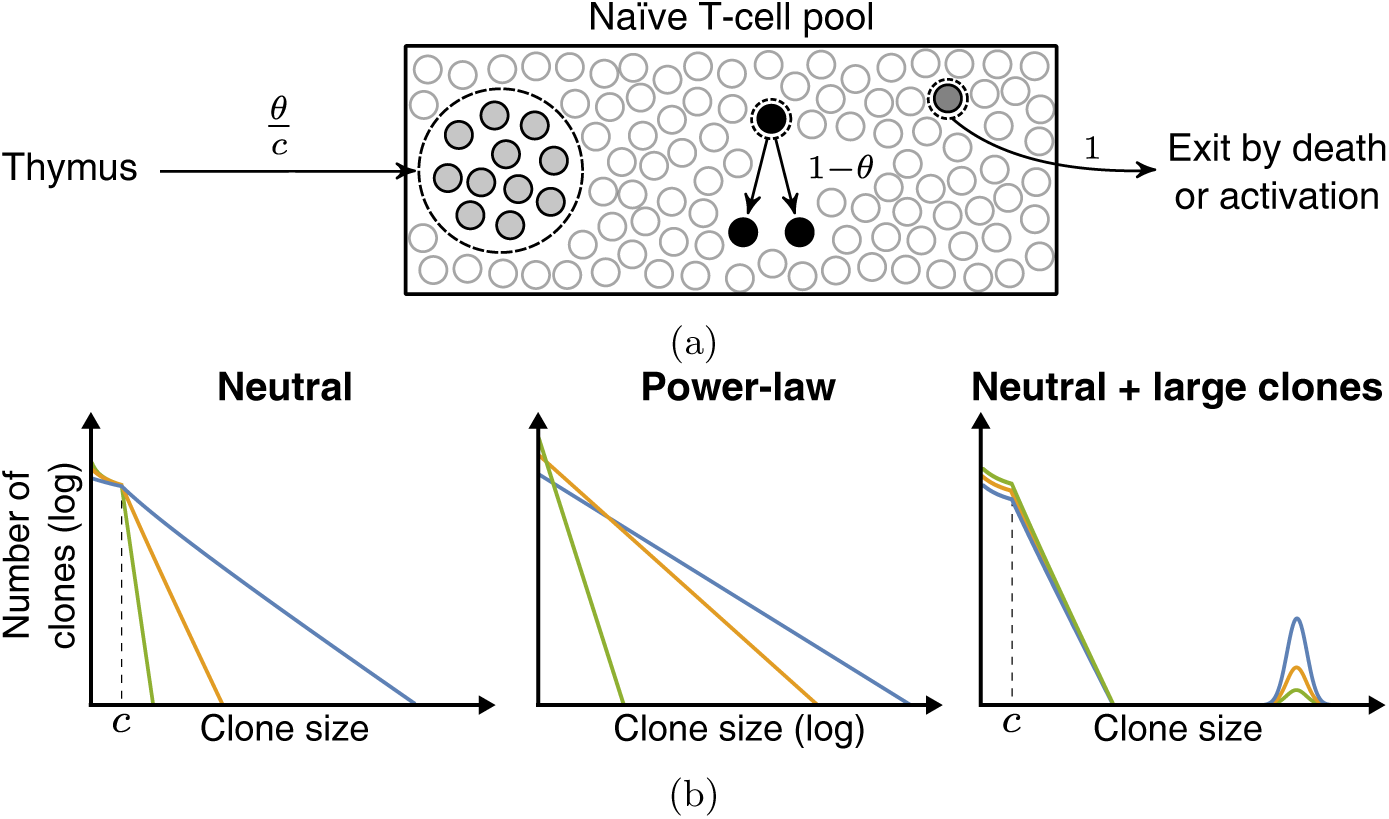
Schematic representation of the neutral model and various clone-size distributions. **A.** Schematic representation of the dynamics of the neutral model for the naive T-cell pool. Each event starts with removal of one randomly selected cell from the pool, followed by peripheral division of another cell (with probability 1 − *θ*), or a chance for thymic production (probability *θ*). After *c* of these thymus events, a clone of *c* cells is generated and added to the peripheral pool, reflecting the divisions of T cells before entering the periphery. **B.** Schematic representations of the various clone-size distributions that were used to predict the naive repertoire. The green, orange and blue colored lines depict three parameter choices for each distribution, resulting in a low, medium and high mean clone-size, respectively.

Samples of immune repertoires are often reported to be power-law distributed [8]. Hence, we compare the predictions of the neutral model with power-law, as well as geometric, log-normal and combined distributions. Note that we vary the shape of each of these phenomenological distributions by changing a single parameter (as shown in Fig. 3B). This allows us to compare distributions with a different degree of heterogeneity in clone sizes. In all cases, we normalize the clone-size distribution such that the total number of cells *N* equals a constant number. To account for the larger CD4^+^ pool [51, 52], we set its pool size *N* = 7. × 5 10^10^ cells, while we used *N* = 2.5 × 10^10^ for the naive CD8^+^ pool.

From all modelled clone-size distributions we take samples and assign *α* and *β* chains that were generated with IGoR [27]. Previous studies showed that *α* and *β* chains with higher generation probabilities tend to have a higher probability to survive selection [11]. Therefore, we train a simple 𝒫(*σ*)-dependent selection model on the data from the single naive T-cell samples shown in Fig. 1. First, we assume that productively rearranged chains have an overall 1*/*3 probability to survive thymic selection. Then we bias the probability for bins of sequences based on their 𝒫(*σ*), such that the resulting set of *α* and *β* chains has the same generation probability distribution as in the HTS data (SI 4.6). Another parameter we learn from the data is the number of cells that contributed at least one mRNA molecule. We set the number of cells that contributed mRNA such that the predicted diversity of a subsample matches the observed diversity (SI 4.6). Taken together, the subsamples we take from the various predicted clone-size distributions are such that they match the generation probabilities and diversity of the HTS subsamples as well as possible. Then we compare the number and 𝒫(*σ*) of the chains that were predicted in multiple subsamples, with those that showed incidence 2 and 3 in the HTS data.

The clone-size distribution which emerges from the neutral model is geometric for clone sizes larger than the introduction size *c* (Fig. 3). As no clone-specific fitness differences are included, the heterogeneity in clone sizes is rather limited, even in case of a minimal contribution of thymic output (*θ*; Fig. 4A). This means that neutral dynamics do not allow clones to become very large, but *α* and *β* chains may be frequent by summation over many of these clones. For the *α* chain, a wide range of thymic output rates *θ* predicts the number of chains occurring in 1, 2 and 3 subsamples reasonably well (Fig. 4A). Although the median 𝒫(*σ*) of predicted incidence 2 and 3 chains does not exactly match the HTS data, the increasing 𝒫(*σ*) with increasing frequency is predicted correctly (Fig. 4D). For the *β* chains, there was no range of thymic output rates that matches the number of incidence 2 and 3 chains simultaneously. Moreover, the observation that incidence 2 chains have higher 𝒫(*σ*) than incidence 3 chains was not predicted for any value of *θ* (Fig. 4D). Thus, the neutral model performs reasonably well in explaining the distribution of *α* chains, but cannot account for the *β* chains in our samples.

**Figure 4:**
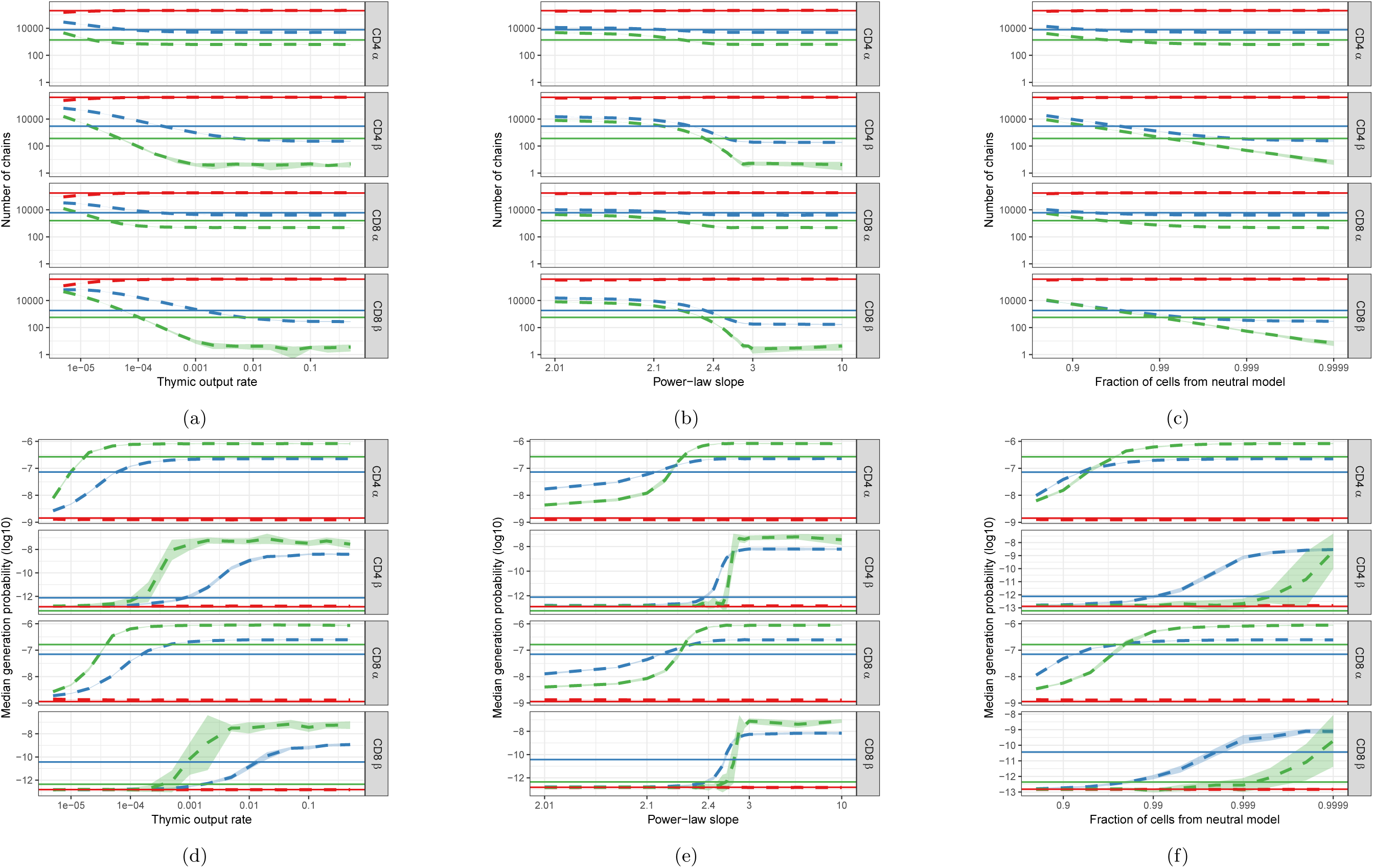
Predictions of the neutral (left), and power-law (middle) and combined model (neutral and large clones, right) compared with HTS data. **A-C.** Number of *α* and *β* chains predicted to occur in 1 (red), 2 (blue) and 3 (green) subsamples as a function of the thymic output rate *θ* for the neutral model (A), the slope of the power-law distribution (B) and the fraction of cells following neutral dynamics in the combined model (C). **D-F.** The median generation probability 𝒫(*σ*) of predicted chains. Dashed lines depict the mean of 10 model prediction repeats, shaded area indicates the standard deviation, solid lines show observed results in HTS data.

### 2.5 The observed distribution of TCR *α* and *β* chains is compatible with an underlying heavy-tailed distribution of T-cell clones in the naive repertoire

We next explored a number of possible empirical frequency distributions for the peripheral naive T-cell repertoire, and modelled the effects of sampling from such populations. As expected from our results with the neutral model, geometric or log-normally distributed T-cell populations were not compatible with the observed data (Fig. S9). Over a wide range of parameters, the distributions account for the number of *α* chains observed in 1, 2 and 3 subsamples, with increasing median 𝒫(*σ*), but fail to predict the number of *β* chains found in 2 and 3 subsamples. Moreover, the predicted median 𝒫(*σ*) of the *β* chains observed in two subsamples was never higher than those with incidence 3. Note that the extreme case in which all clones only consist of one cell, and hence frequently observed chains are the result of summation only, yields similar results. Thus, based on the TCRA data we cannot distinguish between the models above, but the TCRB data shows that the log-normal or geometric distributions of *αβ* clones cannot account for the experimental observations.

We then tested the model using power-law clone-size distributions. Like the distributions discussed before, in the parameter range where clone-size heterogeneity is limited (i.e. a steep slope), a power-law distribution accounts for the number of *α* chains found in 2 and 3 samples, with increasing 𝒫(*σ*). The number of *β* chains with incidence 2 and 3 is predicted well if the slope is close to 2.3 (Fig. 4B). Remarkably, for this slope the predicted incidence 2 *β* chains also showed higher 𝒫(*σ*) than incidence 3 chains (Fig. 4E). Intuitively, we can understand this observation as reflecting the properties of power-law distributions, which contain large numbers of rare events (small clones) but an over-representation of large clones. Sampling from such distributions, there exist parameter ranges in which incidence 3 chains are mostly the result of large clones, while the high 𝒫(*σ*) chains do sum to frequencies that are high enough to be observed in two, but not three subsamples. In such cases, the model would in fact predict the paradoxical observation that the most frequently observed chains have lower 𝒫(*σ*) than less frequent chains.

Finally, we investigated if the agreement between our experimental data and the model is exclusively a property of power-law distributions. We combined the distribution of the neutral model with a variable fraction of large clones (sizes were drawn from a log-normal distribution with clones in the order of 10^5^ cells, SI 4.5). When the large clones make up only a small fraction of the naive T-cell pool (∼1%) the predicted samples match the HTS data as well as those from power-law distributions (Fig. 4C&F). This shows that the explanation of our data does not rely on the specific shape of the power-law distribution. Rather, we find that distributions which contain many small and some very large *αβ* clones are compatible with our TCR sequencing data.

## 3 Discussion

To study the clone-size distribution of TCRs in the naive T-cell pool, we sequenced *TCRA* and *TCRB* mRNA from fractionated blood mononuclear cells, collected from healthy volunteers. Interestingly, sequences frequently observed in the naive repertoire tended to have a higher generation probability 𝒫(*σ*) than less frequent sequences, a trend that is observed much more strongly for TCRA than TCRB sequences. Using a subsampling approach, we confirmed that these trends were not caused by differential mRNA content but by the frequency of *α* and *β* chains in our samples. Surprisingly, the most frequently observed *β* chains appeared to have little or no enrichment for high 𝒫(*σ*). We reasoned that frequent chains can be derived from large clones or alternatively from many clones expressing the same *α* chain but different *β* chains, or vice versa. Our mathematical models for the naive T-cell clone-size distribution explicitly take this summation effect into account, as well as the differential production probabilities of chains and the sampling process. Note that our approach did not focus on finding one unique clone-size distribution that agrees with the experimental data. Instead, we studied a wide range of hypothetical distributions and checked which of these could account for the patterns in our HTS data. We find that the striking observation that TCRA, but not TCRB, frequency can be explained by differential 𝒫(*σ*) is evidence for a broad distribution of clone sizes, i.e. a combination of many small but also a few very large clones.

Our results raise the question which mechanisms allow a small fraction of naive clones to expand to > 10^5^ cells. The neutral model already excluded repeated thymic production as explanation for large clones, because the combined probability of repeated *αβ*-clone production is very low (< 10^−12^ [10]). An alternative explanation is that the large clones may not be truly naive. Although a strict FACS-gating strategy was followed, a small fraction of the cells sorted as naive will be contaminated with (large) memory clones. Alternatively, they could be derived from stem cell memory T cells [18] or other antigen-experienced T cells with a naive phenotype [14, 24, 25, 28, 38]. However, in both these scenarios we would expect cells of these clones to occur also in the memory samples. Excluding all naive sequences that were also observed in corresponding memory samples did not qualitatively change our results (Fig. 2, Fig. S8). We also confirmed that the abundant chains were not strongly enriched for sequences characteristic of iNKT and MAIT cells (SI 4.2). So, although some of the frequent chains may be derived from clones that are not truly naive, our findings still suggest the existence of truly naive large clones. Hence, another possibility is that some peripheral selection (increased survival or proliferation [42]) causes a small number of clones to grow very large. For example, high affinity for self-pMHC is associated with higher fitness [47, 20]. Note that a clone size of 10^5^ cells is theoretically reached after log2(10^5^) ∼17 rounds of divisions. A division rate to achieve this number during a human lifespan does not seem impossible, although the question remains if naive T cells can divide this many times without losing their naive phenotype.

A limitation of the current study is that the experimental setup did not allow for direct analysis of *αβ*- clones. This would be possible by using new techniques that allow for the sequencing of single cells. Current single cell techniques, however, are still limited by sequencing depth. Even though our results indicate that some naive clones are large (> 10^5^ cells), typical sample sizes are too small to observe all of them. One can use the analytical solution of the neutral model (SI 4.4) with thymic introduction size *c* = 1 to illustrate the extreme sampling effect: 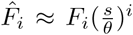, where 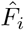 and *F*_*i*_ are the number of clones present with *i* cells in the sample, and in the pool, respectively, and *s* is the fraction of the repertoire that was sampled (here *s* ∼ 10^−6^). Since *s/θ* is of order 10^−5^ and this is raised to the *i*^th^ power, even very large TCR clones become rare in a sample. This shows how difficult it is to infer the full clone-size distribution of the TCR repertoire from small samples, as different distributions tend to converge to samples with a very similar shape.

Our analysis highlights that repeated production of single *α* or *β* chains leads to their occurrence in many different *αβ*-clones. As recombination statistics appear to be similar between people [27], repeated production high 𝒫(*σ*) chains may also account for public sequences. Indeed, sequences which are found in multiple individuals are to a large extent explained by generation probabilities [12]. However, this does not imply the presence of public clonotypes, since there is no reason that the second TCR chain in such clones should also be public. It is necessary to be careful in imputing functional significance to such public TCRA or TCRB sequences, since they probably do not represent public clonotypes. In conclusion, our study provides strong experimental evidence that the human peripheral naive T-cell repertoire contains clonotypes with a broad range of frequencies. This has important functional consequences, since previous studies have shown that the size of the naive pool may determine the strength of the immune response [32]. The mechanisms of peripheral selection which give rise to these distributions remain poorly understood and merit further study.

## Acknowledgements

We thank Laurens Krah for mathematical advice and helpful discussions. This work was supported by The Netherlands Organization for Scientific Research (NWO) Graduate Program 22.005.023 (to P.C.d.G.), the VIRGO consortium, which is funded by the Netherlands Genomics Initiative and by the Dutch government (FES0908) (to B.G.), by a grant to B.C. from Unilever PLC and supported by the National Institute for Health Research UCL Hospitals Biomedical Research. J.H. was supported by an MRC studentship.

## 4 Supplementary Information

### 4.1 Cell sorting and sequencing

Sequence reads came from T cells extracted from blood samples of three healthy volunteers, between 30 and 40 years old. Using CD27 and CD45RA markers, FACS sorting was performed, identifying naïve (CD27^+^CD45RA^+^), CM (central memory, CD27^+^CD45RA^-^), EM (effector memory, CD27^-^CD45RA^-^) and EMRA (effector memory RA CD27^-^CD45RA^+^) cells. Barcoded TCRA and TCRB cDNA libraries were obtained by reverse transcription of RNA molecules coding for either the *α* or *β* chain, respectively, followed by single strand DNA ligation to attach unique molecular identifiers (UMIs) of 12 nucleotides. These were PCR-amplified and sequenced using the Illumina MiSeq platform. For full description of the sequencing procedure, we refer to Oakes *et al.* 2017 [37] and Uddin *et al.* 2019 [50]. The raw sequence files are available on the Sequence Read Archive (https://www.ncbi.nlm.nih.gov/sra) as experiment SRP109035.

### 4.2 Sequence analysis

We used the Decombinator pipeline [49] (Version 3.1) to demultiplex, annotate, and error-correct the raw sequencing reads. Our reads contain UMIs of 12 base pairs that can be used to identify which TCRA or TCRB sequences are derived from the same cDNA molecule. Decombinator performs error correction on sequences by collapsing those that are similar and are associated with the same UMI. The pipeline also error corrects UMIs, collapsing those UMIs that are associated with the same TCRA or TCRB sequence and differ from each other by 2 or fewer sequence edits (i.e. the default barcode threshold). This error correction assumes it is unlikely for any sequence, irrespective of its frequency, to contain two UMIs that are nearly identical, concluding the UMIs are different because of PCR or sequencing errors.

We improved this by setting the barcode threshold to 0 and replacing it by an UMI error correction algorithm that takes the number of UMIs into account. Consider a TCRA or TCRB sequence supported by *i* different UMIs, i.e. with frequency *i*. The Hamming distance, *H*, between two random UMIs of 12 base pairs can be represented by a binomial random variable, *H* ∼ B (*n, p*), where *n* = 12 and 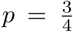 (assuming uniform frequencies of the 4 different bases). There are 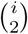 distinct comparisons between the *i* UMIs, and assuming that every comparison is independent, the expected distribution of Hamming distances is 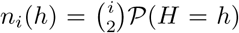. To determine whether two UMIs are unexpectedly similar, we define a threshold distance that depends on the frequency of their TCRA or TCRB sequence (*i*):

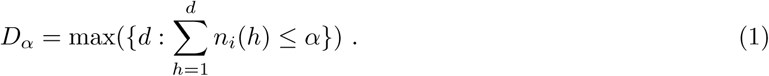

Our algorithm corrects UMIs for a given sequence as follows: From *d* = 1 to *d* = *D*_*α*_, for all UMI pairs with *H* ≤ *d*, add the read count of the less frequent UMI to the more frequent UMI and remove the former. We applied this algorithm to every TCRA and TCRB sequence in our HTS data using *α* = 0.05. The effects of this correction method are shown in Fig. S1. After the improved correction, the distribution of Hamming Distances within and between distinct TCRA and TCRB sequences is very similar, indicating that most erroneous UMIs have been removed. Our improved correction decreases the estimated frequency of many sequences at low frequencies, which indicates that many TCRA and TCRB sequences that were observed two or three times, are actually singletons for which the UMI was mutated once or a few times. In the example given in Fig. S1, the number of sequences that were observed more than once decreased with 66% by our improved correction (from 11491 to 3855), whereas the default correction estimated 9342 (only 19% reduction) of the sequences to have more than 1 true UMI.

**Figure S1:**
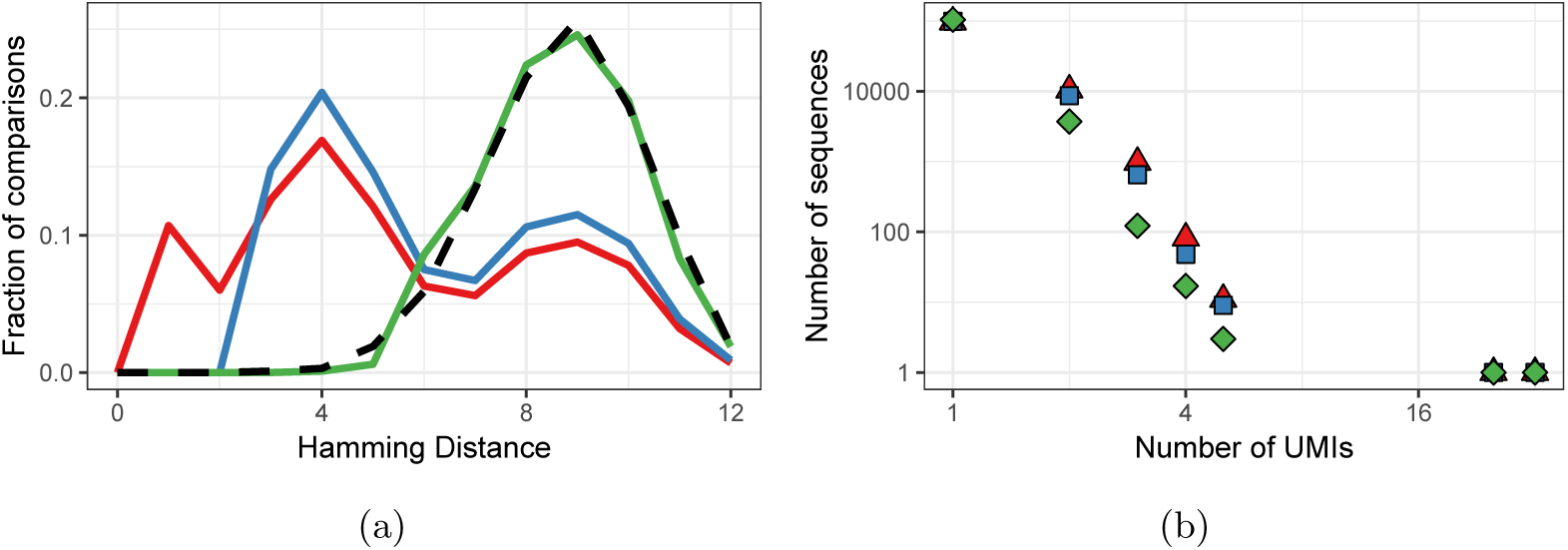
Improved UMI correction leads to more reliable estimation of sequence frequencies. **A.** Distribution of Hamming Distances of UMIs within TCRB sequences (naive CD4^+^ sample of volunteer 1) before correction (red), after default correction (blue) and after improved correction (green), in comparison with the distribution of UMIs between sequences (black dashed). **B.** Distributions of the same TCRB sequences after the different correction strategies. Frequently observed TCRB sequences remain at the same frequency after correction, whereas the frequency of other sequences tends to be overestimated due to mutated UMIs, which is compensated for by improved UMI correction.

We also processed the HTS reads with RTCR [15] (Version 0.4.3). This pipeline determines a sample-based error rate and uses this rate to perform clustering on reads. Compared to Decombinator, RTCR estimates our reads to contain more PCR and sequencing errors and therefore tends to be more conservative in terms of reported diversity. Because RTCR reports fewer distinct rearrangements per sample, the overlap between samples (i.e., the number of chains with incidence 2 and 3) is lower than in Decombinator output. Fig. S5, Fig. S6 and Fig. S7 show the RTCR-based versions of the Fig. 1, Fig. 2 and Fig. 4, respectively. Although the quantitative results are not identical, the RTCR results qualitatively match those of the Decombinator output, confirming that our results are not pipeline-dependent.

Even though a strict FACS-sorting strategy was followed, the CD27^+^CD45RA^+^-sorted sample is expected to contain some (abundant) T cells from the memory compartments. To enrich for truly naive T-cell sequences, one could therefore decide to remove from the analysis any TCRA or TCRB sequence that was also observed in the corresponding memory (CM, EM and/or EMRA) datasets. However, the results shown in Fig. 1A indicate that this cleaning method introduces large biases regarding the 𝒫(*σ*) distribution of the sequences. Indeed, not only chains caused by contamination with very abundant memory clones are removed by this approach, but also *α* and *β* chains occurring in multiple clones (due to high 𝒫(*σ*)). We therefore performed the HTS data analysis on both the original and cleaned data sets and compared them in Fig. 2. In the modelling sections, we proceeded with the original data (including sequences that occurred in memory samples). The results of modelling the cleaned data are shown in Fig. S8. The qualitative agreement between original and cleaned data indicates that our results are not dependent on the removal of possible contamination.

We also checked if the abundant sequences in our data showed characteristics of semi-invariant NKT and MAIT cell populations. Classical NKT cells are characterized by an invariant TRAV24-TRAJ18 *α* chain and *β* chains with TRBV11 [5]. MAIT cells are enriched for TCRA rearrangements of TRAV1-2 with TRAJ33, TRAJ12 and TRAJ20 [40], and TCRB sequences predominantly using TRBV20 and TRBV6 [23]. Since our HTS data does not contain information on *αβ* pairing, we studied both chains separately. Regarding *β* chains, we find that a substantial fraction of the observed TCRB sequences matches the listed characteristics of MAIT cells, and NKT to a lesser extent (Fig. S2). For both cell types, however, this fraction does show a clear relation to incidence, which likely reflects general TRBV usage rather than enrichment for MAIT or NKT cells among abundant sequences. The most abundant TCRA sequences seem to be enriched for NKT characteristics, but still account for only a small fraction of the observed sequences (0.3% and 1.7% for CD4 and CD8, respectively, Fig. S2). Hence, we conclude that only a small fraction of the abundant chains may be derived from clones with a MAIT or NKT cell phenotype and most sequences are abundant for another reason.

**Figure S2:**
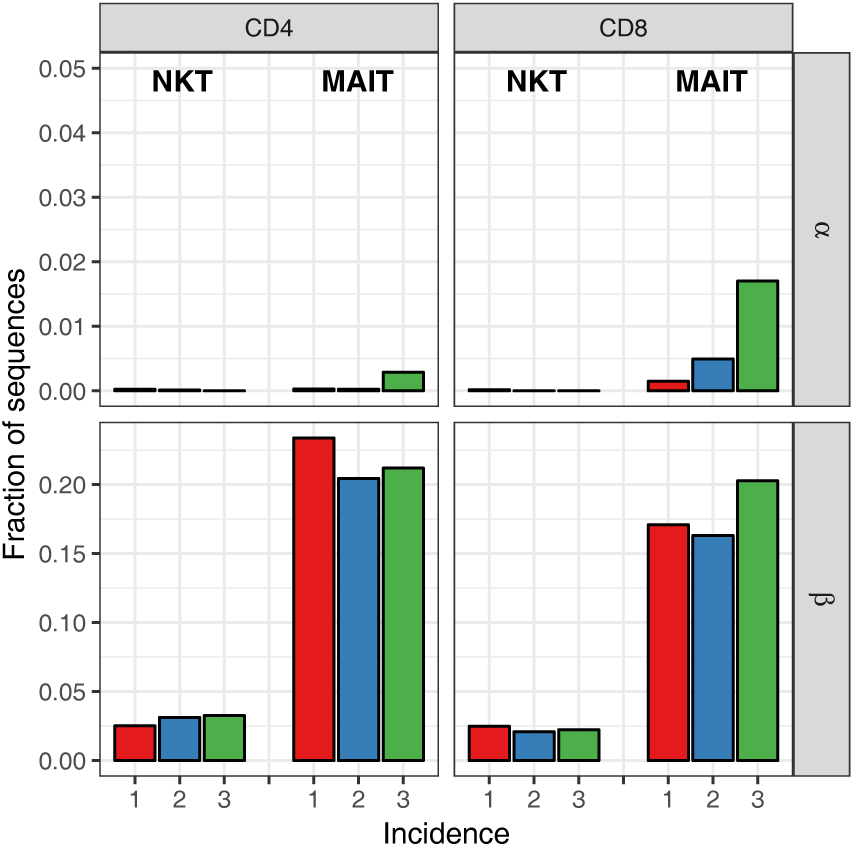
Most abundant chains are not derived from NKT and MAIT clones. Fraction of sequences with rearrangements characteristic for NKT and MAIT phenotypes (as in text).

### 4.3 Sharing of TCRA and TCRB sequences

We sequenced TCRA and TCRB from whole blood samples taken from 28 healthy volunteers. The study was carried out in accordance with the recommendations of the UK Research Ethics Committee with written informed consent of all subjects. All subjects gave written informed consent in accordance with the Declaration of Helsinki. The protocol was approved by the University College London Hospital Ethics Committee 06/Q0502/92. The raw sequence files are available on the Short Read Archive (https://www.ncbi.nlm.nih.gov/sra) as experiments SRP045430 and SRP151125. In order to measure how public the individual sets of sequences were, we measured their degree of sharing between our naive samples and these whole blood repertoires. As shown in Fig. 2, we have three sets of sequences, those with incidence 1, 2 and 3. For each set, we measured which fraction is also found in the 28 independent whole blood samples, which delivers 28 estimates of sharing. More precisely, we counted the number of shared TCRA and TCRA sequences between the sets of sequences observed in two and three naive subsamples, and compared these to sharing with an equal size sample of naive sequences which were only observed in one subsample. Since the number of sequences which occurred more than once was much smaller than the number of sequences which only occurred once, we subsampled the set of unique sequences 10 times. The results are shown as the number of shared TCRA or TCRB for each whole blood repertoire, as a proportion of their number of sequences in the samples being tested (Fig. S3A). In order to study the sharing of the *β* chains in our data with higher resolution, we also analyzed overlap of the sets of sequences with the TCRB data from a large cohort of 786 people published in [13]. The results in Fig. S3B show that a smaller fraction of the most frequently observed *β* chains (incidence 3) are shared than those with incidence 2, which is in line with the P(sigma) observations using IGoR.

### 4.4 Neutral model for dynamics of naive T cells

To model naive T-cell dynamics in the absence of peripheral selection, we developed a model that is similar to the Neutral Community Model (NCM) of Hubbell [19]. Naive T cells, viewed through an ecological lens, are individuals, and all naive T cells sharing the same TCRA and TCRB are part of the same species (*αβ*- clone). Neutrality, as defined by Hubbell, means that all species have the same per capita probability of birth (peripheral division) and death. When considering the model, we ignore the very small chance that an existing *αβ*-clone is produced again by the thymus. Hence, in our simulations we assume that produces T-cell clones that are unique and novel.

**Figure S3:**
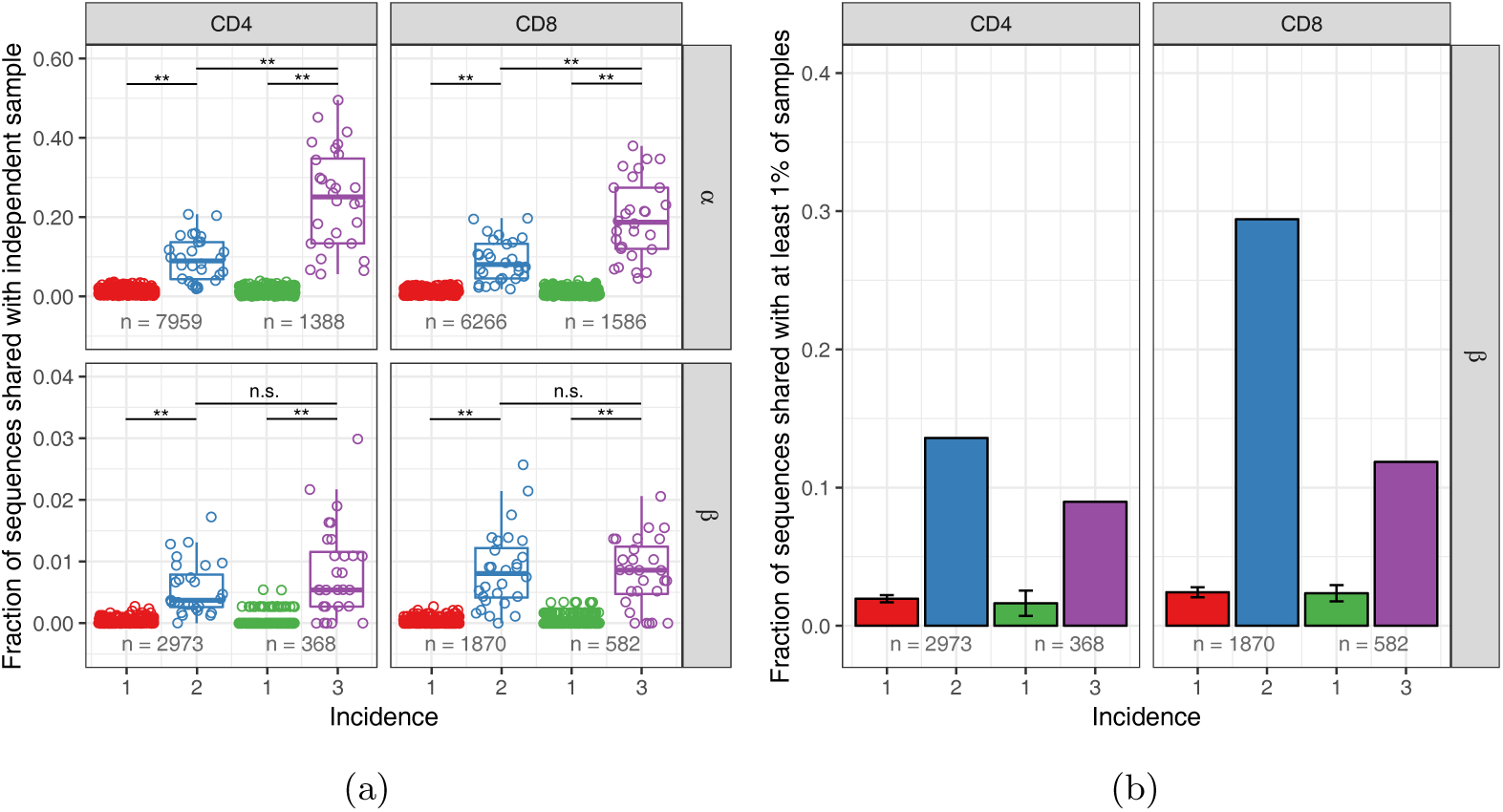
Observed frequency predicts sharing for TCRA but not TCRB sequences. **A.** We compared the occurrence of TCRA and TCRB sequences observed in two or three subsamples (incidence 2 or 3, respectively), and equally-sized samples of the sequences observed in one subsample (incidence 1), in unfractionated blood samples collected from 28 healthy donors. Symbols depict the number of shared TCRA or TCRB sequences for each whole blood repertoire, as a proportion of the total number in the samples being tested (the latter is indicated at the bottom). The boxplot depicts the median value and 25th and 75th percentiles. Shared fractions were compared by Wilcoxon-Mann-Whitney test, **: *p* < 0.01, n.s.: not significant (*p* > 0.05). **B.** Fraction of each set of sequences from A that was observed in at least 1% of the samples from a large cohort of 786 individuals [13]. Error bars show the standard deviation for the multiple sets of sequences with incidence 1.

Consider a pool of *N* naive T cells belonging to clones, each consisting of *i* cells, which changes by thymic production, cell division and cells leaving the naive pool (as a result of cell death or activation). During each event, one randomly selected cell exits the pool, causing the corresponding clone to decrease in size from *i* to *i* − 1 cells. With probability 1 − *θ*, another randomly selected cell will divide, causing the corresponding clone to increase its size from *i* to *i* + 1 cells. Alternatively, with probability *θ*, thymic production *can* occur: every *c* events in which no peripheral division occurred, the thymus will release *c* cells of a newly produced clone. So, the pool size *N* only fluctuates by *c* cells, and because *N* ≫ *c*, the total number of cells stays almost constant during the entire simulation. The per capita birth rate ((1 − *θ*)*/N*) and death rate (1*/N*) are equal for all T-cell clones, which makes this a neutral model. In this discrete-time model, exit and production are coupled, but its dynamics can be approximated by a continuous-time model, in which thymic production, cell division, and deaths are uncoupled Poisson processes. This is illustrated by the following Markov chain, in which the states are clone sizes and the rates show the probabilities of clones moving to another state:

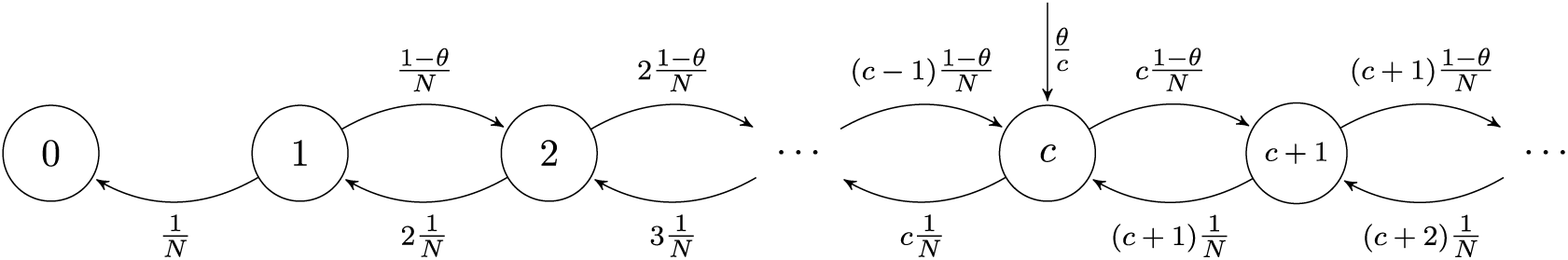

This Markov process describes the dynamics of the clone-size distribution *F*, i.e., the total number of clones *F*_*i*_ consisting of *i* cells. After many birth and death events, individual clones still change in clone size over time, but the clone-size distribution approaches equilibrium. At this steady state, the rate at which new clones enter the naive pool, *θ/c*, equals the rate at which clones leave the pool, i.e., *F*_1_(1*/N*). Hence, in equilibrium, the number of singletons, clones with only one cell, approaches *F*_1_ = *θN/c*. The total rate at which the cells of clones with *i* cells divide and die depends on the total number of cells belonging to *F*_*i*_ clones: *iF*_*i*_. For clone sizes up to *c* cells, the rate at which the cells of the *F*_*i*_ clones die, (*iF*_*i*_*/N*), balances the division the cells of *F*_*i*−1_ clones ((*i* − 1)*F*_*i*−1_(1 − *θ*)*/N*) and the rate at which new clones enter the pool (*θ/c*). The analytical solution to this recurrence relation *iF*_*i*_*/N* = (*i* − 1)*F*_*i*−1_(1 − *θ*)*/N* + *θ/c* is:

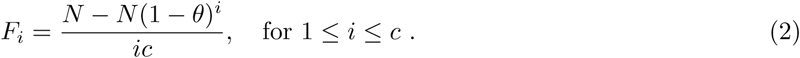

For states with *i* > *c*, only birth and death of cells need to balance between states *i* − 1 and *i* (as there is no net flux from clones introduced by the thymus): *iF*_*i*_*/N* = (*i* − 1)*F*_*i*−1_(1 − *θ*)*/N*. This recurrence relation has the following analytical solution:

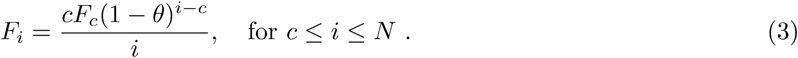

When predicting the full clone-size distribution, we use Eq. 2 and 3 to calculate the steady-state distribution. The total number of all distinct clones (i.e. the richness) in the steady-state repertoire is simply the sum over all their frequencies 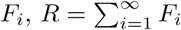, which has a simple closed-form solution for *c* = 1,

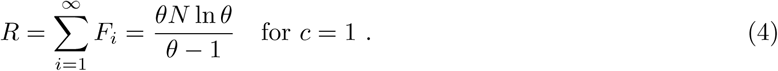

The Simpson’s diversity of the steady state repertoire also has a simple form,

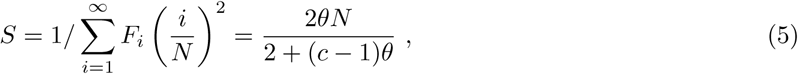

which equals *F*_1_ = *θN* for *c* = 1, and is a saturated function of *θ* if *c* > 1.

We consider the sampling process of a small fraction *s* from a naive T-cell pool of *N* cells, which clones follow the distribution *F* in Eq. 2 and Eq. 3. Assuming the naive pool is large and well-mixed, the number of T cells, *X*, sampled from the *j* cells belonging to a particular clone, can be approximately represented by a binomial random variable, *X*_*j*_ ∼ B (*n* = *j, p* = *s*). The expected clone-size distribution of the sample, 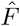, is then given by

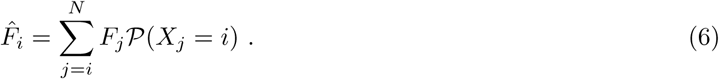

The strong distortion of sampling from clone-size distributions can be illustrated using the analytical solution of Eq. 6 for the neutral model for *c* = 1:

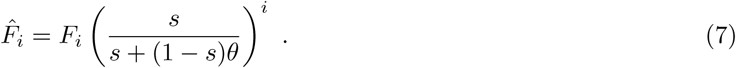

Since *s* is typically very small, this equation can be simplified to 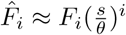 (as *s* ≪ *θ*), which clearly shows that even very abundant clones will become rare or absent in a small sample.

### 4.5 Clone-size distributions of the naive T-cell pools

Since our data contains separate data on both CD4^+^ and CD8^+^ T cells, we predicted the clone-size distributions of both subsets separately. To account for the larger CD4^+^ pool [51, 52], we set its pool size *N* = 7.5 × 10^10^ cells, while we used *N* = 2.5 × 10^10^ for the naive CD8^+^ pool. When analyzing the neutral model, we used its steady-state distribution (Eq. 2 and Eq. 3). Since the *β* chain rearranges first, followed by a few divisions before rearrangement of the *α* chain [16], we use *c* = 100 for TCRB and *c* = 10 for TCRA. We also used various phenomenological clone-size distributions that are not based on a mechanistic model. To allow for exploration of a wide range of distributions, we chose mathematical functions which form can be changed by a single parameter, such as the slope of the power-law and geometric distribution. The power-law distribution with form *F*_*i*_ = *F*_1_ × *i*^−*k*^ shows a straight line on a log-log plot. Since all *F*_*i*_ are written as a function of *F*_1_, the total number of cells 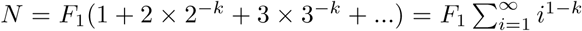. This sum is convergent for *k* > 2 and gives

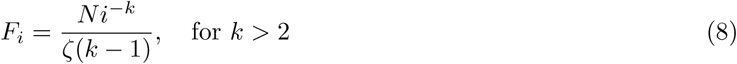

for the power-law clone-size distribution, in which *ζ* is the Riemann zeta function. The geometric distribution, with form *F*_*i*_ = *F*_0_ × *b*^*i*^ is a straight line on a semi-log plot. Requiring 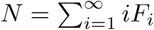 yields

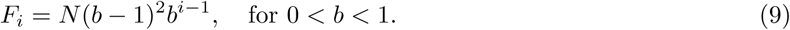

We also studied repertoires with log-normal distributions of clone-sizes by drawing from a normal distribution and raising 10 to the power of these numbers for clone sizes. For this we used varying *µ* and *σ* = *µ/*10. These distributions yielded results that were qualitatively similar to those from the geometric distribution (Fig. S9). Lastly, we combined the aforementioned distribution following from the neutral model with a population of large clones. We define a fraction *f* for cells following the distribution of the neutral model using the Eq. 2 and 3 with *N* ^*\**^ = *f N*. We then added the remaining (1 − *f*)*N* cells following the approach for the log-normal distribution, with *µ* = 5.

### 4.6 *In silico* samples from modelled clone-size distributions

To compare the clone-size distributions with the HTS-data of the blood samples, we generated TCRA and TCRB repertoires using IGoR [27]. We generated 10^8^ TCRA and TCRB sequences using IGoR’s default recombination model and parameters. We selected the rearrangements which CDR3 nucleotide sequence consisted of a multiple of 3 nucleotides (in-frame) and did not contain in-frame stop codons, in line with the inclusion criteria of productive rearrangements in our HTS samples (∼28%). Next, we calculated generation probabilities 𝒫(*σ*) for all these rearrangements. This may seem a detour, but this is needed as many different scenarios can lead to the same TCRA or TCRB rearrangement.

Only a small percentage of thymocytes that undergo rearrangements in the thymus will eventually be exported as a naive T cell. This is due to out-of-frame rearrangements, but also as a result of both positive and negative selection. Moreover, the generation probability distributions of pre- and post-selection TCRA and TCRB repertoires are markedly different [11]. To account for these observations, we train a 𝒫(*σ*)- dependent selection model to account for the effects of thymic selection on our IGoR-produced TCRA and TCRB sequences. Note that this selection method is based on single chains rather than *αβ*-TCRs. We do this because the 𝒫(*σ*)-shift shows that selection does happen on single chains (i.e., an *α*-chain can be selected against irrespective of the *β*-chain or vice versa). Most likely selection also acts on the level of *α* and *β* chains together, but randomly removing combinations of these would not alter the eventual distribution of the single chains that are observed in our data.

We use each of the HTS data sets from the single sample experiment (shown in Fig. 1) to calculate the relative enrichment or depletion of 100 log10 𝒫(*σ*) bins (ranging from −50 to 0) compared to 100 equally sized samples of the IGoR output, for TCRA and TCRB separately. If the HTS data contained few rearrangements for a given bin, we joined adjacent bins in such a way that the bin-specific selection factor was always based on at least 1% of the experimental observations (Fig. S4). This approach yielded 𝒫(*σ*)-specific selection factors *f*_𝒫(*σ*)_ ranging from 0.6 to 1.15 (i.e., our data suggests that sequences with a preferable 𝒫(*σ*) are about 2 times as likely to be selected as those in the least preferable 𝒫(*σ*) domain). We assumed an overall selection factor *f*_*overall*_ of 1*/*3, meaning that one out of 3 productive TCRA and TCRB rearrangements would survive selection. We then allowed sequences to be part of the post-selection repertoire with probability

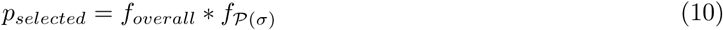

and stored the outcome to make a consistent decision when multiple copies of the same TCRA or TCRB sequence were present in the pre-selection repertoire. This approach yielded a post-selection repertoires with 𝒫(*σ*) distributions similar to the single sample HTS data. Other values of *f*_*overall*_, ranging from 1*/*10 to 1 were also tested, but yielded similar qualitative results (not shown).

**Figure S4:**
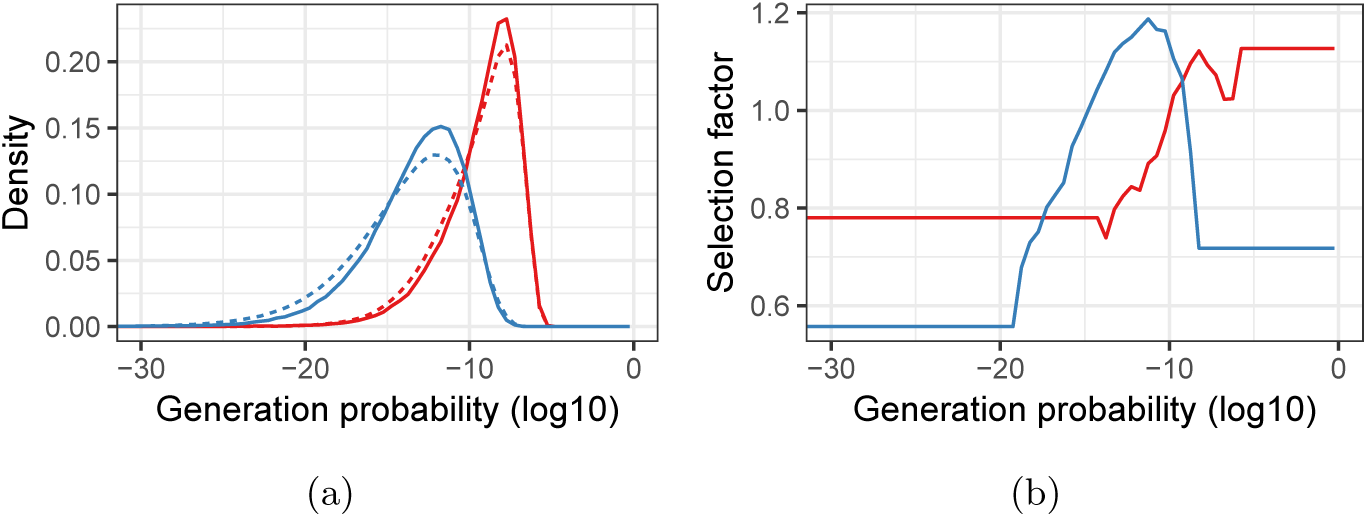
Pre- and post-selection 𝒫(*σ*) densities and 𝒫(*σ*)-dependent selection factors for *α* and *β* chains. **A.** Relative frequency of generation probabilities of TCRA (red) and TCRB (blue) sequences in the combined HTS data (solid) and IGoR output (dashed). **B.** The bin-specific selection factors *f*_𝒫(*σ*)_ are determined by division of the density of a given bin in the HTS data by the density in the pre-selection IgoR output. A value of 1 means that a sequence with this 𝒫(*σ*) has an average probability to be selected in the thymus, whereas lower values indicate stronger selection and higher values weaker selection (i.e., a higher probability to pass selection).

We could have assigned all clones in the clone-size distribution an *α* and *β* chain with this approach. However, since only a very small part of the repertoire is sampled, we chose to only assign an identity to those clones present in the samples. Hence, we started with predicting the presence of all clones, as a function of their size, in each of the samples. The probability that a clone with *i* cells is represented by at least one cell in a sample of *n* cells from a pool of *N* cells is

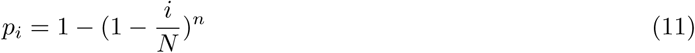

Given *F*_*i*_, which is the number of clones in the pool with clone size *i*, the number of these clones present in the sample of *n* cells can be approximately represented by a binomial random variable, *X*_*i*_ ∼ *B*(*n* = *F*_*i*_, *p* = *p*_*i*_). We evaluate this for the entire clone-size distribution *F*. *N* and *F* are known from the model but one cannot directly determine the number of sampled cells *n*. This is because individual cells may contribute multiple mRNA molecules and many cells may have been present in the FACS sorted sample without contributing mRNA to the eventual sequenced fraction. Therefore we learn the sample size by assigning *α* or *β* to sampled clones and choosing *n* such that the predicted diversity (i.e., number of distinct chains) matches the experimental observations. We took the number of distinct TCRA or TCRB sequences as lower bound for the sample size, since in this model individual cells are assumed to express one functional *α* or *β* chain. The total number of cells reported by the FACS-sorter was used as upper bound. We also checked the implications of the observation that some T cells contain two functional *α* and/or *β* chains, but this did not qualitatively change our results (not shown).

Thus, we adjusted the generation probability distribution by training a 𝒫(*σ*)-dependent selection model on independent HTS data and based the sample size on the corresponding subsamples. Hence, the predicted individual subsamples reflect the experimental observations in terms of diversity and generation probabilities. We use the chains occurring in multiple samples (i.e., those with incidence 2 and 3) to assess the agreement between model predictions and the HTS data. We repeated the sampling process and assignment of *α* and *β* chains 10 times for each model-parameter combination to account for the stochastic nature of sampling and V(D)J recombination.

**Figure S5:**
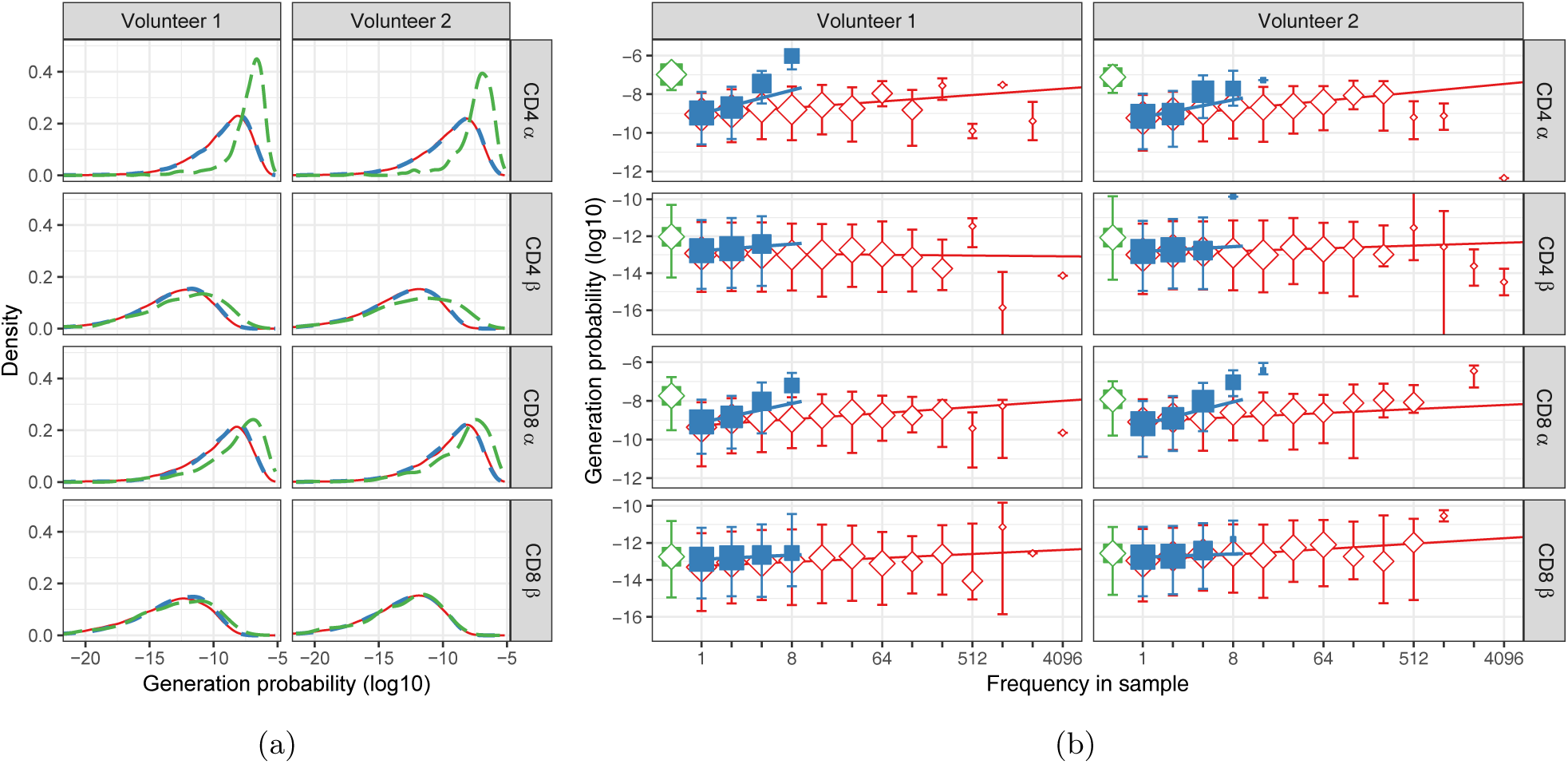
Similar to Fig. 1, but for HTS data processed with RTCR. **A.** For each sequence *σ* in our dataset, the generation probability 𝒫(*σ*) was predicted using IGoR [27]. The distribution of generation probabilities (log10) for *α* and *β* chains from CD4^+^ and CD8^+^ from two volunteers is shown. Blue dashed: naive, red solid: memory, and green long-dashed: overlap (i.e., sequences observed in both naive and memory). **B.** The median 𝒫(*σ*) is shown for each observed frequency class (log2-bins) in naive (blue squares) and memory T-cell (red diamonds) samples. 𝒫(*σ*) of the overlapping chains is shown in green for reference (irrespective of frequency). Symbol sizes indicate numbers of sequences for each frequency class. Error bars represent the 25% and 75% quartiles, solid lines indicate linear regression between observed frequency and 𝒫(*σ*), weighted by the number of sequences with that frequency.

**Figure S6:**
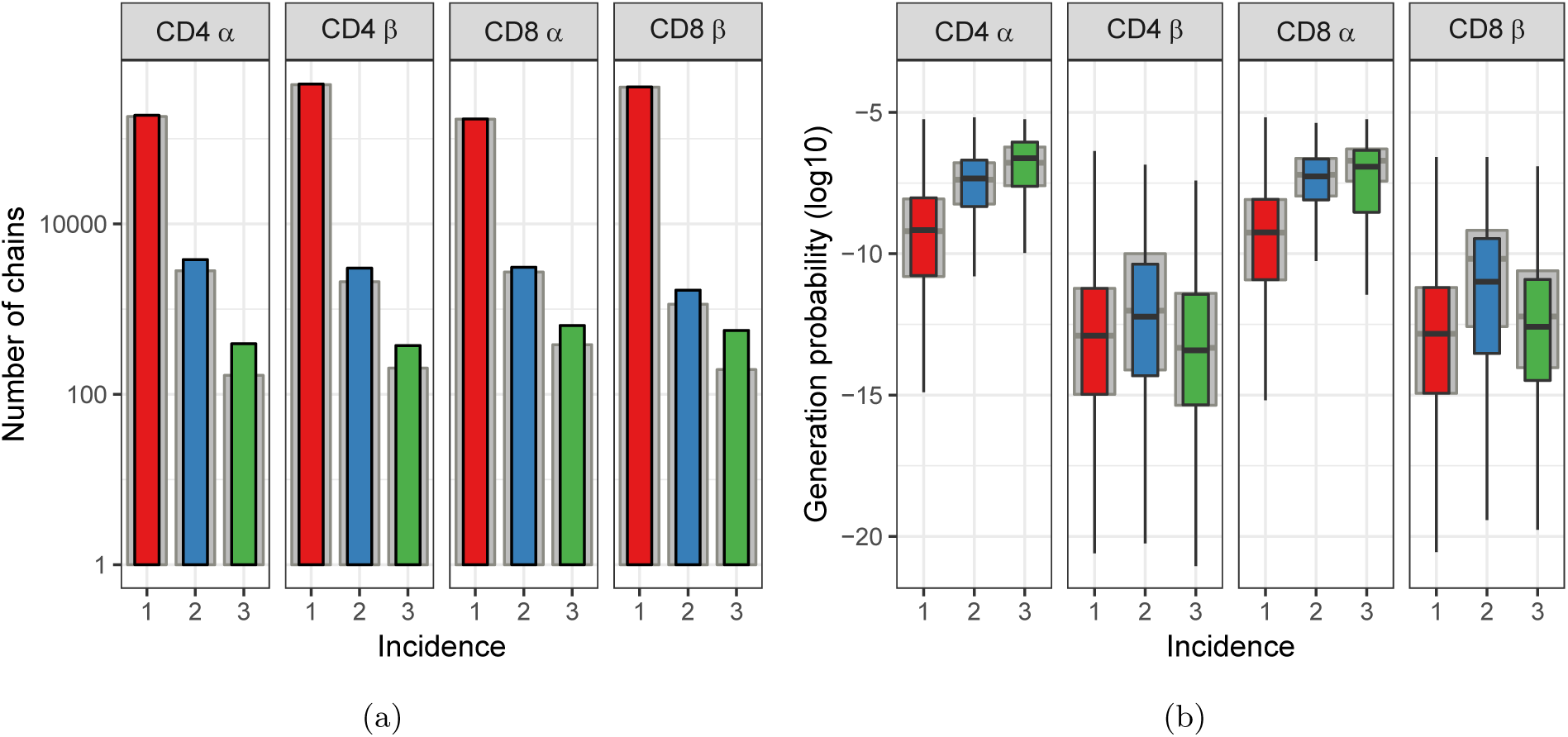
Similar to Fig. 2, but for HTS data processed with RTCR. **A.** The number of TCRA and TCRB sequences observed in 1, 2 or 3 subsamples. The grey background bars show the results after removing all sequences that were also observed in the corresponding memory samples. **B.** Generation probabilities 𝒫(*σ*) (log10) of chains observed in 1, 2 or 3 subsamples.

**Figure S7:**
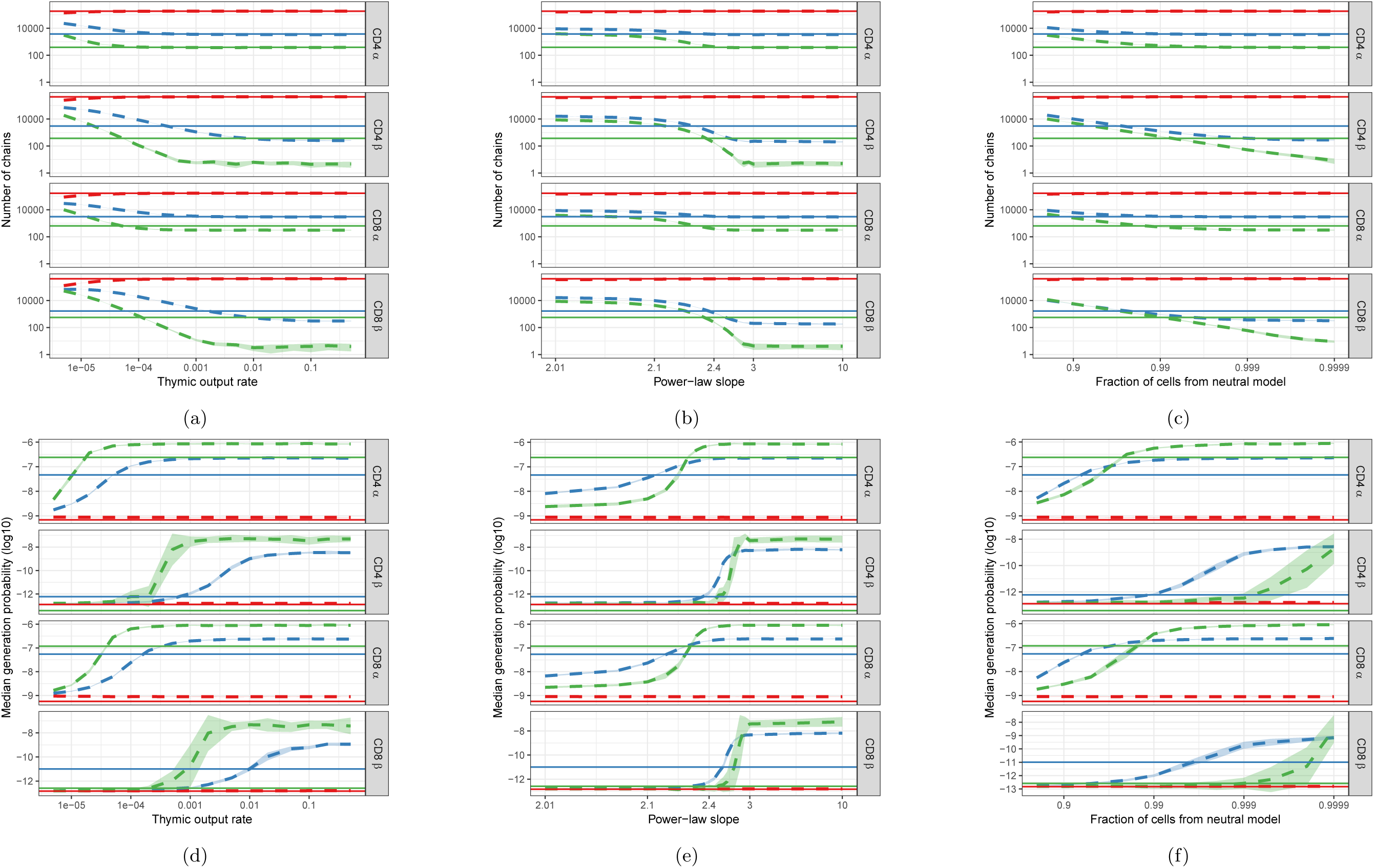
Similar to Fig. 4, but for HTS data processed with RTCR. **A-C.** Number of *α* and *β* chains predicted to occur in 1 (red), 2 (blue) and 3 (green) subsamples as a function of the thymic output rate *θ* for the neutral model (left), the slope of the power-law distribution (middle) and the fraction of cells following neutral dynamics in the combined model (right). **D-F.** The median generation probability 𝒫(*σ*) of predicted chains. Dashed lines depict the mean of 10 model prediction repeats, shaded area indicates the standard deviation, solid lines show observed results in HTS data.

**Figure S8:**
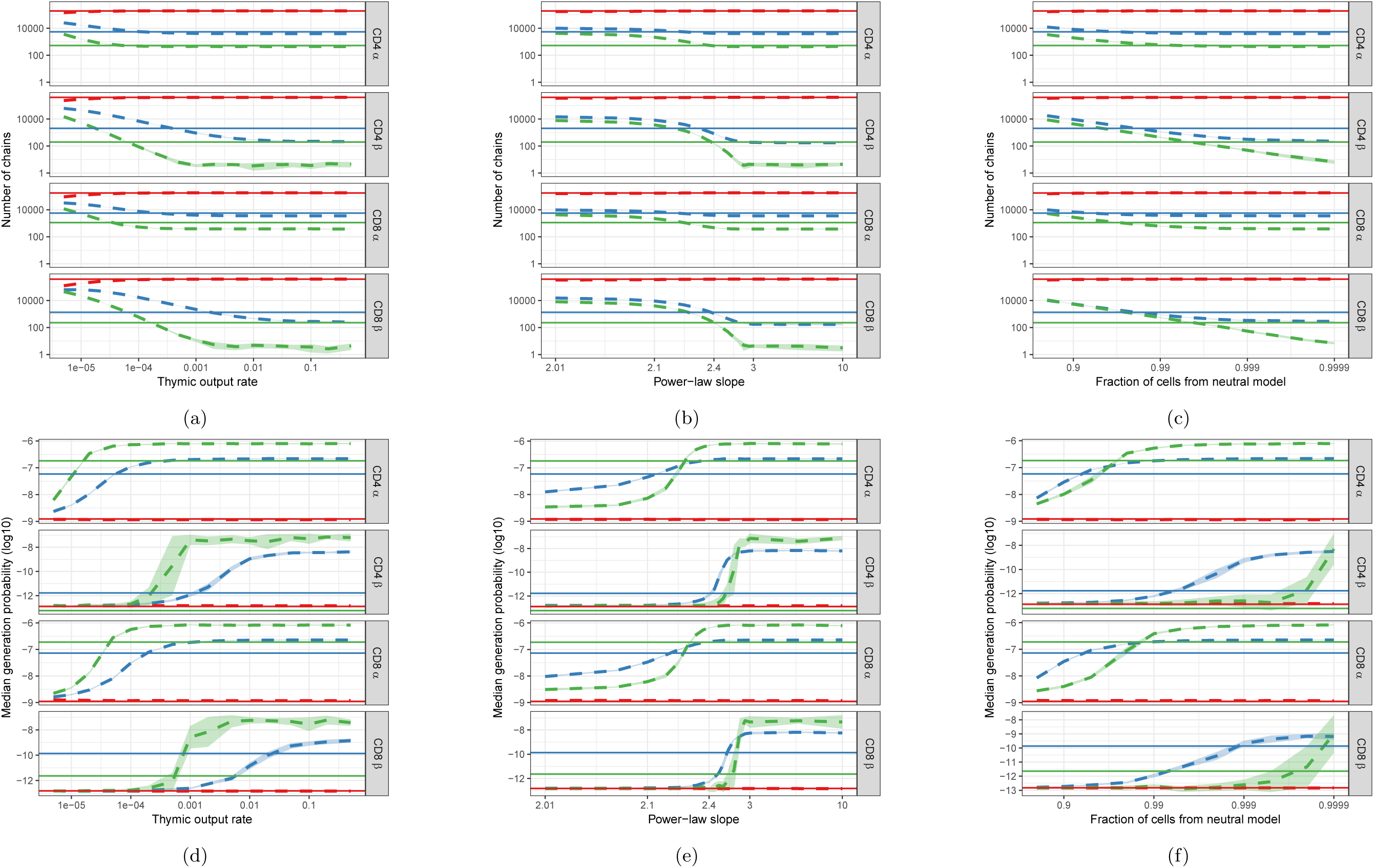
Similar to Fig. 4, but for HTS data from which TCRA and TCRB sequences were removed that also occurred in the corresponding memory samples. **A-C.** Number of *α* and *β* chains predicted to occur in 1 (red), 2 (blue) and 3 (green) subsamples as a function of the thymic output rate *θ* for the neutral model (left), the slope of the power-law distribution (middle) and the fraction of cells following neutral dynamics in the combined model (right). **D-F.** The median generation probability 𝒫(*σ*) of predicted chains. Dashed lines depict the mean of 10 model prediction repeats, shaded area indicates the standard deviation, solid lines show observed results in HTS data.

**Figure S9:**
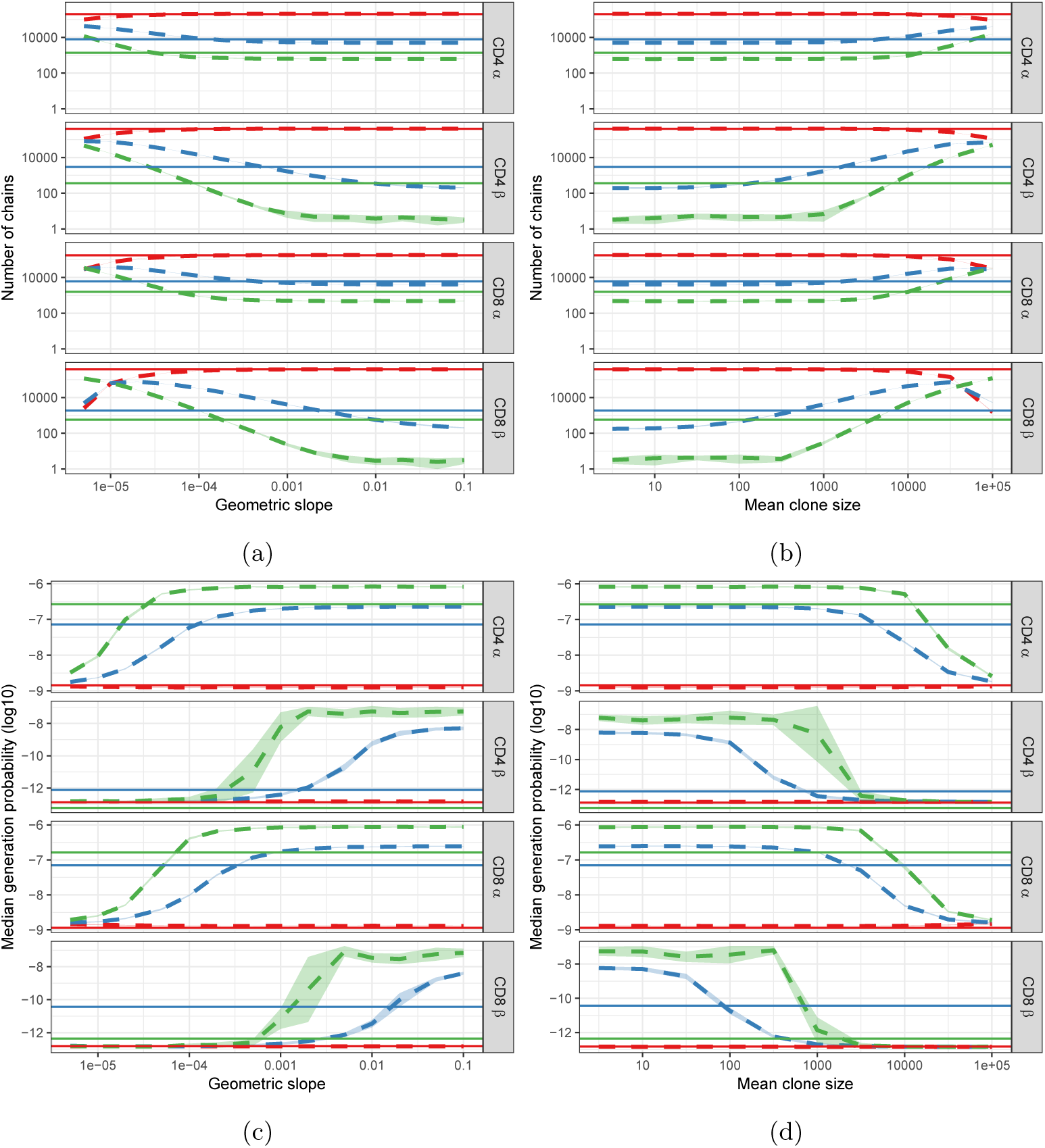
Predictions of the geometric (left) and lognormal (right) distribution compared with HTS data. **A&B.** Number of *α* and *β* chains predicted to occur in 1 (red), 2 (blue) and 3 (green) subsamples as a function of the slope *b* for the geometric distribution (left) and mean clone size for the lognormal distribution (right). **C&D.** The median generation probability 𝒫(*σ*) of predicted chains. Dashed lines depict the mean of 10 model prediction repeats, shaded area indicates the standard deviation, solid lines show observed results in HTS data.

